# Hypothalamic CRH neurons gate rapid social appraisal of conspecifics

**DOI:** 10.1101/2025.10.31.685874

**Authors:** Ibukun Akinrinade, Meenakshi Pardasani, Toni-Lee Sterley, Tamás Füzesi, Jaideep S. Bains

**Author notes:** Equal contribution (Ibukun Akinrinade, Meenakshi Pardasani).

## Abstract

Successful navigation of the social environment requires rapid evaluation of conspecifics to distinguish safe from threatening encounters. While the corticotropin-releasing hormone neurons in the paraventricular nucleus of the hypothalamus (CRH^PVN^) are well known for orchestrating endocrine stress responses, their role in real-time social appraisal is unknown. Here, using fibre photometry and optogenetics during ethologically relevant social interactions, we show that CRH^PVN^ neurons exhibit a rapid, transient increase in activity that precedes sniffing, an investigative behaviour through which mice gather social information. This transient signal scales with social context: it is heightened during interactions with unfamiliar and aggressive conspecifics and significantly reduced toward familiar partners. Inhibiting these neurons selectively diminished anogenital sniffing, demonstrating their necessity for optimal social evaluation. Strikingly, across repeated exposures to the same conspecific, behavioural investigation declined while CRH^PVN^ activity remained robust, revealing a dissociation between neuronal and behavioural habituation. This persistence suggests that CRH^PVN^ neurons encode stable neural representation of threat during social encounters. Together, our findings identify CRH^PVN^ neurons as a fast, integrative node linking endocrine stress control to rapid social decision-making.

## INTRODUCTION

Individuals are constantly exposed to various threats in their environment. Their survival critically relies on their ability to rapidly assess and evaluate such threats. While threat detection has been studied extensively in the context of predator-prey interactions^1–3^, less is known about how individuals navigate the more subtle but equally consequential risks posed by conspecifics. Encountering an unfamiliar individual prompts a series of behaviours designed to gather information in order to assessment potential risk^4^. Although interactions with conspecifics can be rewarding, fostering development, reproduction, and emotional resilience^5,6^, they can also pose a threat, leading to aggressive interactions, competition for limited resources and establishment of dominance and aggressive interactions which promote a stress response^7,8^. Importantly, social threats are often ambiguous and uncertain, requiring optimal speed and accuracy in detection. The delayed detection of an actual threat can result in severe outcomes such as injury or death, while overreacting to a non-threat may incur unnecessary social and metabolic costs^9,10^. How individuals evaluate their conspecifics and the neural substrates underlying these behaviours are poorly understood.

CRH^PVN^ neurons, the canonical controllers of the endocrine response to stress, have emerged as key regulators of threat-related information^11–17^. They are necessary for the social investigation of stressed partners, a behaviour that facilitates the transmission of stress between individuals^14^. Additionally, these neurons show increased activity in response to social stimuli^18^. Despite these findings, the role of CRH^PVN^ neurons in gauging the threat posed by their conspecifics is unclear. Here, we use an ethologically relevant social behaviour paradigm, combined with real-time neuronal activity measurements and optogenetics, to show the direct involvement of CRH^PVN^ neurons in gating social threat behaviours. Our data show that a rapid increase in CRH^PVN^ neuron activity is required for social investigation, a behaviour necessary for the optimal assessment of conspecifics. Together, these findings identify CRH^PVN^ neurons as critical components of the neural circuitry underlying social appraisal.

## RESULTS

### Context-dependent investigative behaviours during real social assessment

To investigate how the threat level posed by a conspecific is determined, we first evaluated behavioural changes during safe and threatening social encounters by exposing mice to familiar cage mates and aggressor strain, respectively (Figure 1A). Directed sniffing (a dynamic respiratory behaviour essential for information gathering^20–23^) initiated by the resident mouse was assessed during the first 3 minutes of social interaction. We analysed four distinct sniffing behaviours (Figure 1B): anogenital sniffing (snout of resident directed toward the anogenital region of intruder), head sniffing (snout of resident directed toward the head of intruder), torso sniffing (snout of resident directed toward the trunk of intruder) and tail sniffing (snout of resident directed toward the tail region of intruder). Analysis of transition probabilities revealed that social investigation patterns differed markedly between familiar and aggressor encounters. Compared with familiar interactions, encounters with aggressors produced much stronger Head→Torso transitions and higher probabilities for the Torso→Anogenital and Anogenital→Torso transitions. This pattern suggests that during aggressor interactions, residents follow a more directed sequence from Head→Torso→Anogenital regions and a more repeated alternation between the torso and anogenital regions. In addition, the Tail→Anogenital transition increased, suggesting that residents rapidly directed their investigation toward the anogenital region when faced with an aggressor (Figure 1C).

**Figure 1:**
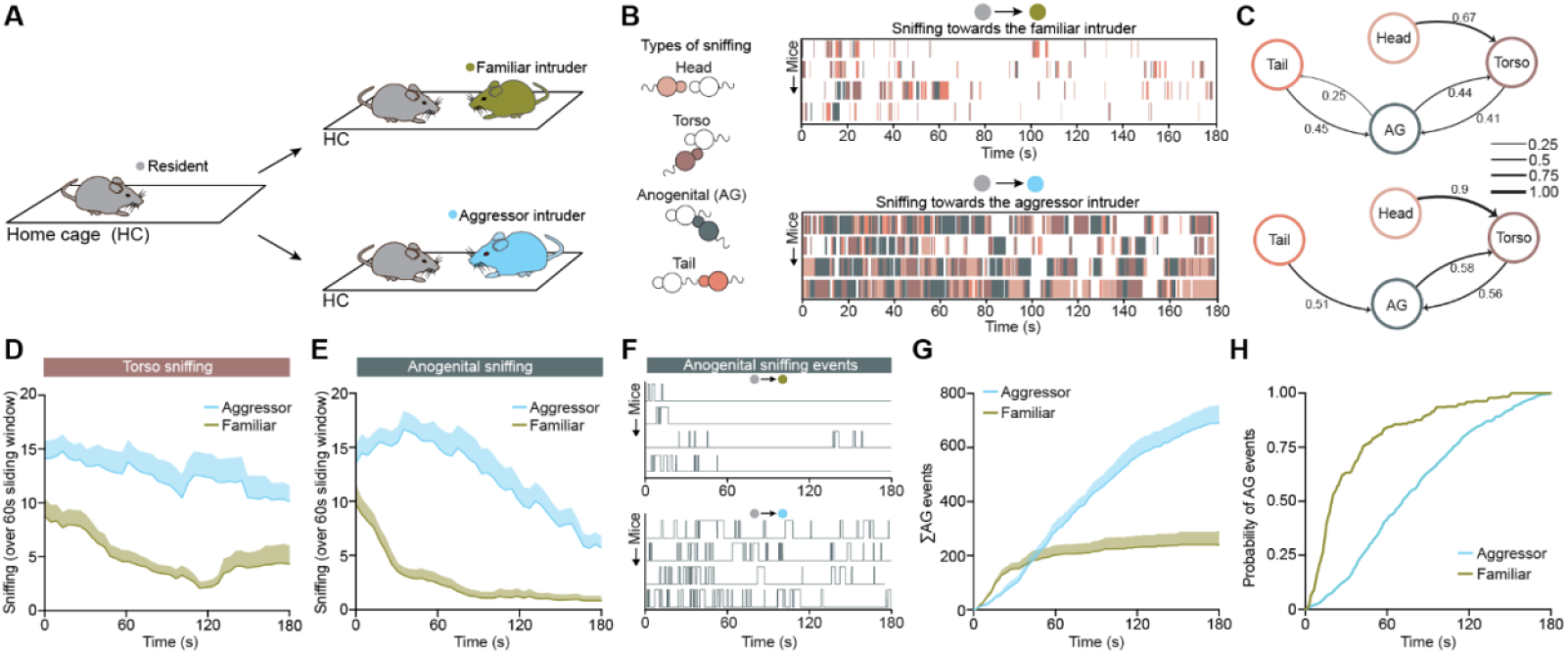
Resident mice exhibit targeted investigation of a threat-inducing intruder. **A**. Schematic of Resident-Intruder test in home cage of a resident mouse interacting with either a familiar conspecific (green) intruder or an aggressor CD1 mouse (blue) intruder. **B**. Example behavioural sequences of investigative sniffing events exhibited by the resident mouse towards the different anatomical positions on the body of the intruder (anogenital, head, torso, and tail). **C**. Transition probability plots representing switching of sniffing behaviours towards familiar (top) and aggressor (bottom) mice for a total of 3 min. **D**. Mean torso sniffing time averaged in 60s bins with a 3s sliding window during the 3 min-long interaction (N = 12 for both groups – 2-way RM ANOVA Group X time: F (41,902) = 0.76, p = 0.855, shade represents s.e.m.). **E**. Mean anogenital sniffing time averaged in 60s bins with a 3s sliding window during the 3 min-long interaction (N = 12 for both groups – 2-way RM ANOVA Group X time: F (41,902) = 4.89, p < 0.0001, shade represents s.e.m.). **F**. On-off dynamics of each anogenital sniffing event. **G**. Group average of cumulative duration for anogenital sniffing (N = 12 for both groups – 2-way RM ANOVA Group X time: F (3000, 66000) = 28.99, p < 0.0001, shade represents s.e.m.). **H**. Cumulative probability plots showing the distribution of anogenital (AG) sniffing events, with curve indicating the cumulative likelihood of AG event occurrence in each social context.

Next, we compared the temporal dynamics of sniffing behaviours across the 3 min interaction period. While torso sniffing followed a similar time course in both familiar and aggressor encounters, anogenital sniffing showed a significant divergence between groups, increasing steadily during aggressor encounters but remaining lower and more transient in familiar interactions (Figure 1D,E). In addition, head sniffing showed a modest temporal variation, with a comparable early time course followed by a slight increase in investigation toward the latter phase. Tail sniffing, however, did not differ significantly between groups, indicating that this behaviour remained stable across social contexts (Figure S1).

Given that aggressor encounters suggest a shift in the pattern of investigation toward the anogenital region, we next focused on a detailed analysis of anogenital sniffing^23–27^. The anogenital region concentrates social chemosignals that convey critical chemosensory information that aids social evaluation^27–30^. Sustained investigation of this region likely reflects an enhanced drive for social threat evaluation amongst other functions. To determine how this behavioural focus evolves over time, we next quantified the temporal dynamics of anogenital sniffing across the 3-min interaction. Raster plots reveal the fine temporal structure of discrete anogenital sniffing bouts, while the cumulative sum and probability visualize how rapidly and persistently each group accumulates investigative events over time. These data show that exposure to an aggressor elicits denser and more sustained anogenital sniffing, consistent with a prolonged state of social threat assessment. Whereas interactions with familiar conspecifics are characterized by brief, transient sampling typical of non-threatening encounters (Figure 1F-H).

### CRH^PVN^ neuron activity increases during social appraisal of an aggressor

Considering that agonistic encounters can be threatening^31,32^, we sought to determine whether CRH^PVN^ neurons are recruited during social appraisal to evaluate the threat level of an intruder. To monitor real-time activity in freely behaving animals, we used transgenic mice expressing GCaMP6f selectively in CRH^PVN^ neurons (Figure 2A). An optical fibre was implanted above the PVN to perform fibre photometry recordings of calcium-dependent fluorescence as a proxy for population activity^33^ in resident mice during social interactions (Figure 2B).

**Figure 2:**
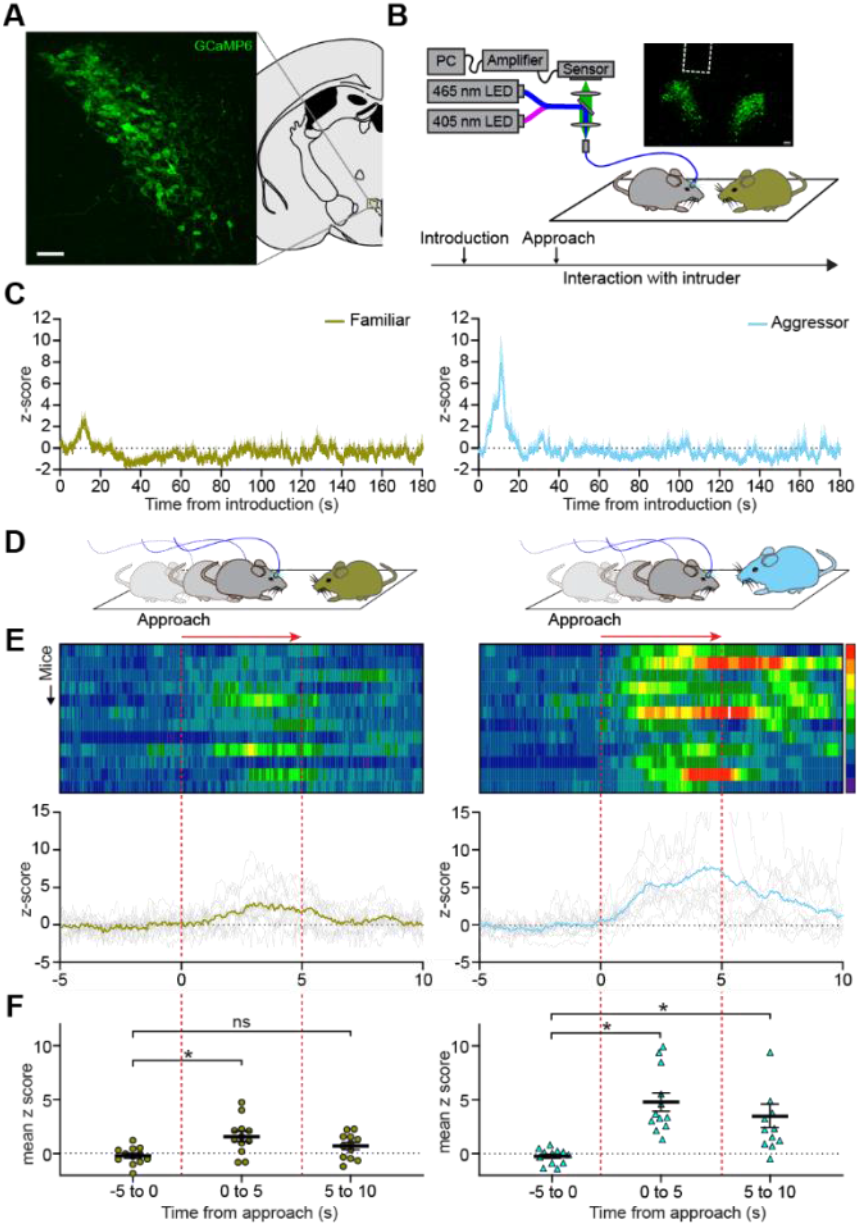
CRH^PVN^ activity increases during social appraisal of an intruder. **A**. Mouse coronal half-section depicting the Paraventricular nucleus (PVN) of the hypothalamus with a confocal image on the left showing GCaMP6f expression in the CRH^PVN^ neurons (scale bar = 50µm). **B**. Schematic depicting fibre photometry from GCaMP6f expressing CRH^PVN^ neurons during social interaction. Confocal image depicting the fibre track above CRH^PVN^ neurons (Right, scale bar = 100µm) and the scheme of social interaction (bottom). **C**. Group average trace of the z-scored calcium activity during social interaction of 3 min towards familiar (left) and aggressor (right) intruder. **D**. Schematic of resident approaching familiar intruder (top), heatmap of calcium activity during approach and corresponding mean z-scored activity graph denoted in green with individual animals depicted in gray (middle), bar graph depicting change in z-score averages across the 5s time windows (w1,w2,w3), with 0s denoting approach (N=12, Repeated measures 1-way ANOVA, F (2,22) = 9.61, p = 0.001 Tukey’s multiple comparison test, w1 vs. w2 p = 0.004, w1 vs w3 p =0.103, w2 vs w3 p = 0.105). **E**. Schematic of resident approaching aggressor intruder (top), heatmap of calcium activity and corresponding mean z-scored activity graph denoted in green with individual animals depicted in gray (middle), bar graph depicting change in z-score average across the 5s time windows, with 0s denoting approach(N=12, Repeated measures 1-way ANOVA, F (2,22) =18.26, p < 0.0001 Tukey’s multiple comparison test, w1 vs. w2 p = 0.0002, w1 vs w3 p =0.009, w2 vs w3 p = 0.227).

The approach by a familiar intruder was accompanied by a transient increase in CRH^PVN^ activity in the resident. Activity returned to baseline within seconds and remained around baseline during the remainder of the 3-minute interaction period. In contrast, the approach toward an aggressor intruder elicited a higher amplitude increase in CRH^PVN^ activity. These initial changes occurred within the first few seconds of intruder introduction in both conditions, indicating a rapid neuronal response to social stimuli (Figure 2C). Closer temporal analysis of each condition revealed that although both familiar and aggressor introductions produced increases CRH^PVN^ activity, the presence of an aggressor triggered a significantly greater response (Figure 2D-F, S2A-D). Together, these findings indicate that CRH^PVN^ neurons are rapidly engaged during social appraisal, with response magnitude reflecting the perceived threat level of the conspecific.

### Unfamiliar conspecific elicits CRH^PVN^ activity that mirrors that evoked by an overt social threat

To determine whether CRH^PVN^ neurons scale to graded levels of social threat, we exposed resident mice to same-strain, same-sex, non-cage mate (unfamiliar) intruders to represent a socially ambiguous, potentially threatening conspecific (Figure 3A). Much like exposure to an aggressor encounter, behavioural ethograms revealed robust investigative sniffing directed toward the unfamiliar intruder, reflecting heightened social appraisal relative to familiar interactions (Figure 3B, S3A-C). Across the 3-minute interaction period, residents spent significantly more time engaged in sniffing behaviours when confronted with an unfamiliar or an aggressive intruder than with a familiar one (Figure 3C, S4A-B). The distribution of sniffing types further revealed an increased proportion of anogenital investigation during both the unfamiliar and aggressor encounters (Figure 3D–E, S4C-D), consistent with the elevated response to potentially threatening social stimuli. Next, we assessed the possible changes in CRH^PVN^ activity in the resident mouse during unfamiliar intruder exposure. Fibre photometry recordings revealed a sharp and transient increase in CRH^PVN^ activity during approach (Figure 3F,H, S5A-B). The magnitude and temporal profile of this activation closely resembled that observed during aggressor encounters (Figure 3G,I,J, S6A-F), suggesting a shared pattern of neuronal activation across contexts of potential and actual social threat. Together, these results indicate that CRH^PVN^ neurons are recruited not only by overtly aggressive encounters but also by exposure to unfamiliar, potentially threatening conspecifics, supporting their role in rapid threat detection.

**Figure 3:**
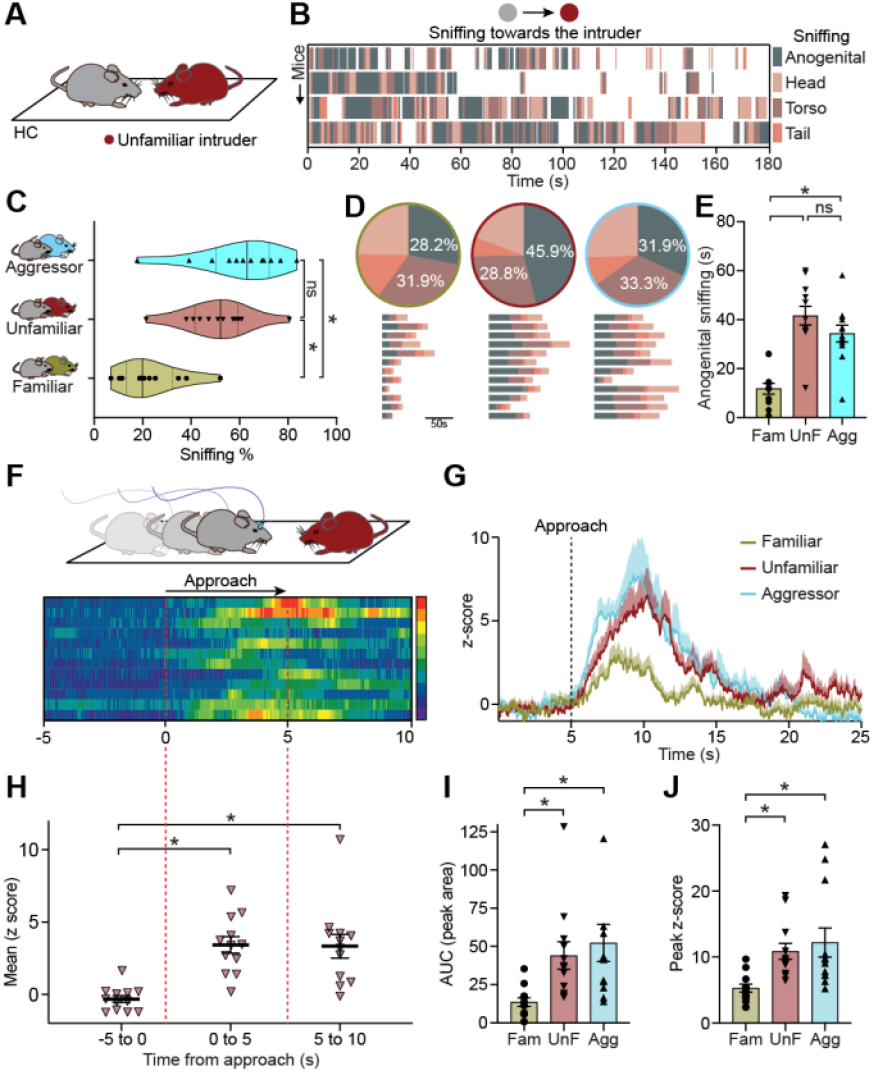
Unfamiliar conspecific intruder increases CRH^PVN^ activity and investigative behavioural display in resident. **A**. Schematic of Resident-Intruder test with an unfamiliar conspecific (maroon) intruder. **B**. Example behavioural sequences of investigative sniffing events exhibited by the resident mouse towards an unfamiliar intruder. **C**. Violin plots depicting percentage of total sniffing by residents during 3-min long social interaction towards the three types of intruders, opaque line represents median and dotted lines represent quartiles q1: 25% and q3: 75% percentiles (N = 12 per group, Ordinary 1-way ANOVA, F(2,33) = 18.6, p < 0.0001, Tukey’s multiple comparison test, Familiar vs. Aggressor p < 0.0001, Familiar vs. Unfamiliar p = 0.0003, and Unfamiliar vs. Aggressor p = 0.343). **D**. Pie charts (top) representing the distribution of sniffing types across familiar, aggressor and unfamiliar intruder groups, and corresponding horizontal bars (bottom) representing time spent in different sniffing events for each animal. **E**. Anogenital sniffing time for 3 groups (data is represented as mean ± s.e.m., N = 12 per group, Ordinary 1-way ANOVA, F (2,33) = 23.15, p < 0.0001, Tukey’s multiple comparison test, Familiar vs. Aggressor p < 0.0001, Familiar vs. Unfamiliar p < 0.0001, and Unfamiliar vs. Aggressor p = 0.263). **F**. Schematic of resident approaching an unfamiliar intruder (top), heatmap of CRH^PVN^ calcium activity and corresponding mean z-scored activity graph denoted in green with individual animals depicted in gray (middle), bar graph depicting change in z-score average across the 5s time windows, with 0s denoting approach(N=12, Repeated measures 1-way ANOVA, F (1.601,17.61) =19.86, p < 0.0001 Tukey’s multiple comparison test, w1 vs. w2 p < 0.0001, w1 vs w3 p =0.002, w2 vs w3 p = 0.98). **G**. Group average z-score of CRH^PVN^ calcium responses aligned at approach, shade represents s.e.m. **H**. Individual peak area under the curve for the calcium activity for the first peak after approach (N = 12 per group, Ordinary 1-way ANOVA, F (2,33) = 5.74, p = 0.0072, Tukey’s multiple comparison test, Familiar vs. Aggressor p = 0.01, Familiar vs. Unfamiliar p = 0.029, and Unfamiliar vs. Aggressor p = 0.90). **I**. Individual peak z-scores of the first peak after approach (N = 12 per group, Ordinary 1-way ANOVA, F (2,33) = 6.08, p = 0.0056, Tukey’s multiple comparison test, Familiar vs. Aggressor p = 0.007, Familiar vs. Unfamiliar p = 0.027, and Unfamiliar vs. Aggressor p = 0.857).

### Neuronal–behavioural dichotomy during repeated social appraisal

Social encounters with unfamiliar or overtly aggressive conspecifics elicit robust investigative and neuronal responses in resident mice. To determine whether these neuronal responses adapt as the social context becomes predictable, we repeatedly exposed resident mice to the same unfamiliar or aggressor intruder across three consecutive days and monitored both social investigation behaviour and transient CRH^PVN^ activity. This analysis aimed to distinguish behavioural habituation from neuronal adaptation, testing whether CRH^PVN^ activity reflects social novelty or a persistent representation of threat salience.

Across repeated encounters in both conditions, resident mice displayed behavioural habituation, characterized by significant reductions in anogenital sniffing, consistent with a decreased investigative drive as familiarity with the intruder increased. In contrast, CRH^PVN^ activity remained robust across sessions, showing no significant decline in event magnitude and duration over days (Figure 4A-D, S7A-F). These findings reveal a dissociation between behavioural and neuronal habituation during repeated social encounters. Thus, CRH^PVN^ neurons appear to provide a stable internal representation of threat that endures even as observable behaviour habituates.

**Figure 4:**
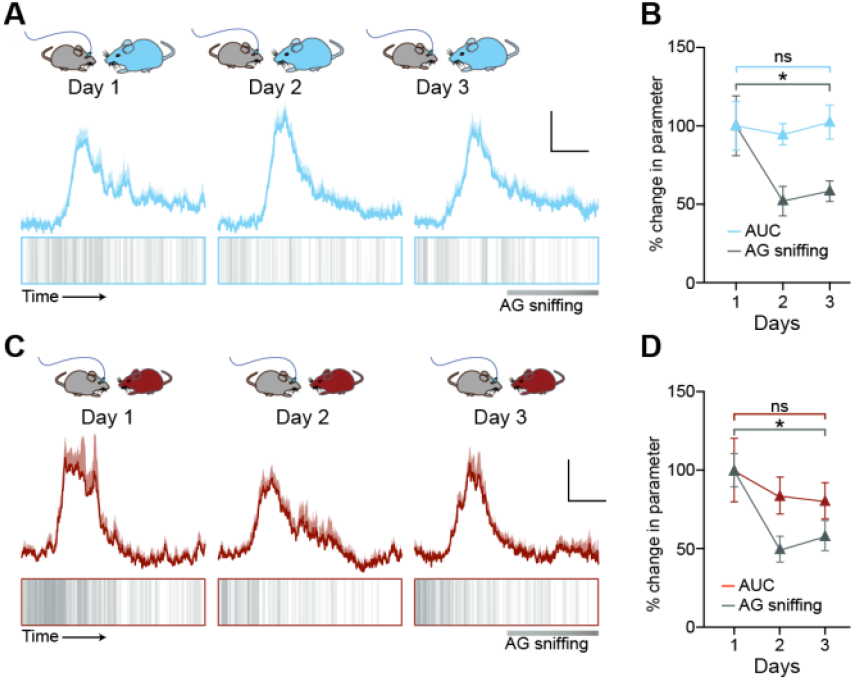
Repeated exposure to unfamiliar and aggressive intruders elicits neuronal-behavioural dichotomy. **A**. Top: Maintenance of group average z-scored CRH^PVN^ calcium responses during interaction with the same aggressive intruder over 3 days of re-exposures, Bottom: anogenital events towards the intruder overlaid for all animals belonging to the same group. **B**. Percentage change in the AUC for the calcium activity in A and anogenital sniffing over 3 days (N =11, 1-way RM ANOVA, AUC: F (1.89,18.99) = 0.1, p = 0.887 and AG sniffing: F (1.31,13.15) = 6.74, p = 0.016). **C**. Top: Maintenance of group average z-scored CRH^PVN^ calcium responses towards identical unfamiliar intruder over 3 days of re-exposures, Bottom: anogenital events towards the unfamiliar intruder overlaid for all animals belonging to the same group. **D**. Percentage change in the AUC for the calcium activity in C and anogenital sniffing over 3 days (N =10, 1-way RM ANOVA, AUC: F (1.29,11.63) = 0.68, p = 0.459 and AG sniffing: F (1.4,12.7) = 14.14, p = 0.0012).

### CRH^PVN^ activity impacts targeted anogenital investigation during social threat appraisal

To determine the necessity of CRH^PVN^ activity neurons in social threat appraisal, we used mice expressing Archaerhodopsin 3.0 in CRH neurons (CRH^Arch3.0^) and optogenetically inhibited^34^ CRH^PVN^ neurons (Figure 5A). During behavioural testing, resident mice received continuous yellow-light illumination for 20s, to suppress CRH^PVN^ activity just as the intruder was introduced (Figure 5B). The timing of inhibition was selected as it aligns with the average period during which the rise and decay of CRH^PVN^ activity occur after exposure to an intruder. Inhibition of CRH^PVN^ activity selectively disrupted sniffing investigative behaviour toward unfamiliar and aggressor intruders without altering investigation of familiar intruders (Figure 5C). Sniffing distributions and event rasters confirmed that photo-inhibition did not alter the duration of anogenital investigation toward familiar mice (Figure 5D-H). Interestingly, inhibition significantly reduced anogenital investigation toward aggressor intruders, indicating a loss of sustained investigative drive in response to threat (Figure 5I-M). Similarly, optogenetic silencing of unfamiliar intruders significantly decreased anogenital investigation (Figure 5N-R). This suggests that rapid activation of CRH^PVN^ neurons is essential for gauging the threat posed by a conspecific and that diminished activity within this population shifts social evaluation toward a neutral state (Figure S8A-E).

**Figure 5:**
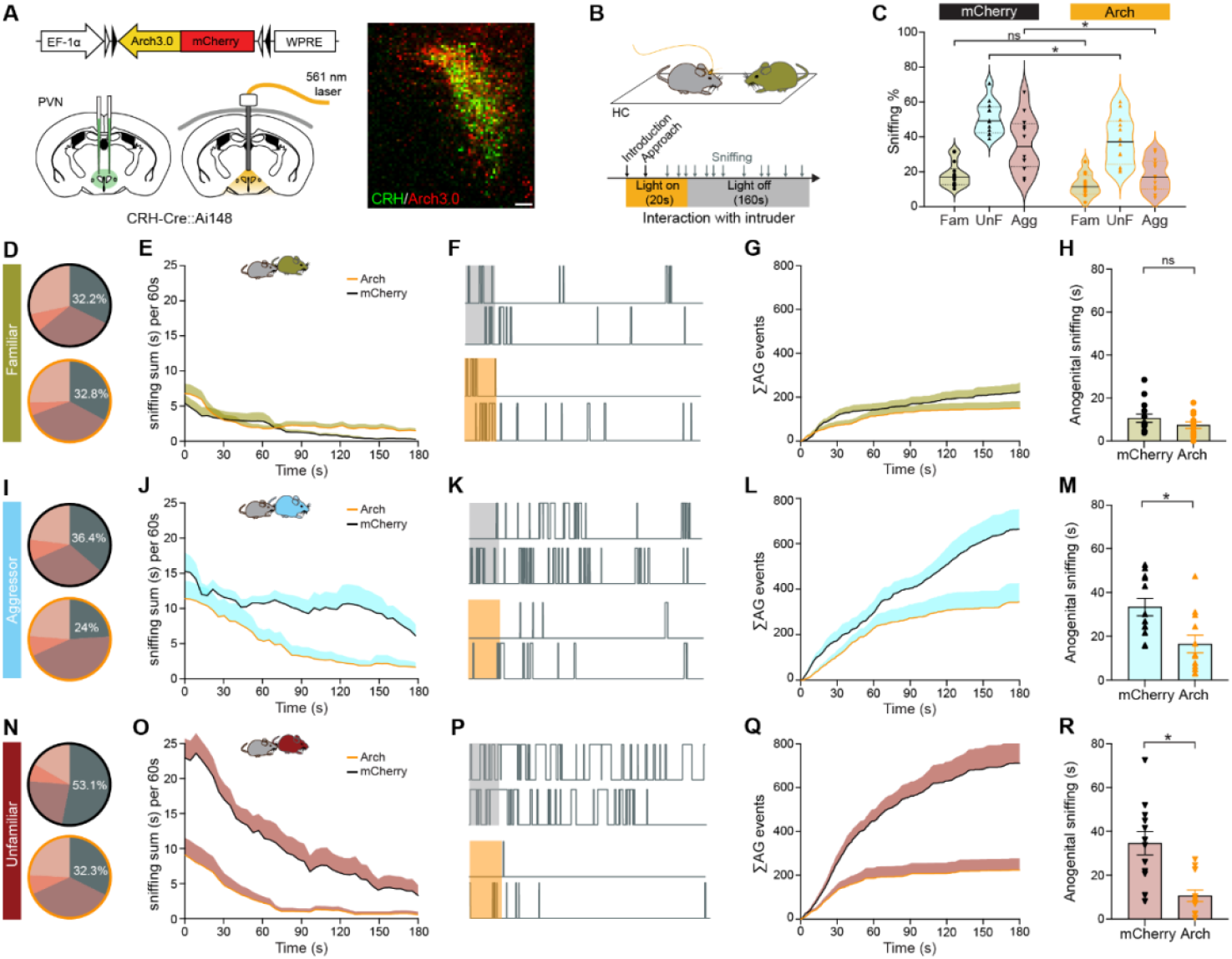
Increase in CRH^PVN^ activity is essential for targeted anogenital investigation of a threat-inducing intruder. **A**. Top: Viral construct for optogenetic inhibition experiment, bottom: Strategy of virus injection and ferrule implantation for optogenetic inhibition of CRH^PVN^ neurons, right: Confocal image depicting Cre-dependent localization of aav2/9-EF1a-DIO-Arch3.0mCherry-CNP virus in the CRH^PVN^ neurons (Scale bar = 50 μm). **B**. Experimental timeline depicting photo-inhibition of CRH^PVN^ neurons in the resident mouse during the first 20s of the social appraisal of an intruder. **C**. Violin plots of the percentage of total sniffing by control (mCherry) and opto-inhibited (arch) residents during 3-minute-long social interaction towards the familiar, aggressor and unfamiliar intruders (N=12-14 per group, 1-way ANOVA, F (5,68) = 23.32, p < 0.0001, Sidak’s multiple comparison test, familiar-arch vs familiar-mCherry p = 0.433, aggressor-arch vs aggressor-mCherry p = 0.015, unfamiliar-arch vs. unfamiliar-mCherry p = 0.0004). **D**. Pie chart representing similar distribution of anogenital sniffing of a familiar intruder between mCherry and arch groups. **E**. Mean anogenital sniffing time averaged in 60s bins with a 3s sliding window during the 3 min-long interaction (N = 12 for arch and 14 for mCherry, – 2-way RM ANOVA Group effect: F (41,984) = 1.38, p = 0.056, shade represents s.e.m.). **F**. Example raster plots of anogenital sniffing events towards familiar intruder (top: mCherry group, bottom: arch group). **G**. Group average of cumulative duration for anogenital sniffing (N = 12 for arch and 14 for mCherry – 2-way ANOVA Group X time: F (3599, 86400) = 0.217, p > 0.99, shade represents s.e.m.). **H**. Bar graph depicting mean ± s.e.m. of anogenital sniffing time for mCherry vs. arch residents towards familiar intruders (N=12 for arch and 14 for mCherry, Unpaired t-test, p = 0.214). **I**. Pie chart representing differences in distribution of anogenital sniffing of an aggressor intruder between mCherry and arch groups. **J**. Mean anogenital sniffing time averaged in 60s bins with a 3s sliding window during the 3 min-long interaction (N = 12 for both groups – 2-way RM ANOVA Group effect: F (41,861) = 1.485, p = 0.027, shade represents s.e.m.). **K**. Example raster plots of anogenital sniffing events towards aggressor intruder (top: mCherry group, bottom: arch group). **L**. Group average of cumulative duration for anogenital sniffing (N = 12 for both groups – 2-way ANOVA Group X time: F (3599, 75600) =1.301, p < 0.0001, shade represents s.e.m.). **M**. Bar graph depicting mean ± s.e.m. of anogenital sniffing time for mCherry vs. arch residents towards aggressor intruders (N=12 for both groups, Unpaired t-test, p =0.007). **N**. Pie chart representing differences in distribution of anogenital sniffing of an unfamiliar intruder between mCherry and arch groups. **O**. Mean anogenital sniffing time averaged in 60s bins with a 3s sliding window during the 3 min-long interaction (N = 12 for both groups – 2-way RM ANOVA Group effect: F (41, 902) =5.53, p < 0.0001, shade represents s.e.m.). **P**. Example raster plots of anogenital sniffing events towards unfamiliar intruder (top: mCherry group, bottom: arch group). **Q**. Group average of cumulative duration for anogenital sniffing (N = 12 for both groups – 2-way ANOVA Group X time: F (3599, 79200) = 3.121, p < 0.0001, shade represents s.e.m.). **R**. Bar graph depicting mean ± s.e.m. of anogenital sniffing time for mCherry vs. arch residents towards unfamiliar intruders (N=12 for both groups, Unpaired t-test, p = 0.0005).

## DISCUSSION

Here, we describe a mechanism that controls the rapid detection of social threat. We demonstrate that the appearance of a conspecific triggers an immediate recruitment of CRH^PVN^ neurons. We also show that inhibiting this activity has a sustained impact on the investigative behaviour required for optimal assessment of the conspecific. By combining detailed behavioural analysis, fibre photometry, and optogenetic inhibition, we identify a functional link between CRH^PVN^ neurons and investigative behaviour. Furthermore, where behavioural responses habituated CRH^PVN^ activity persisted, revealing a neuronal substrate for social vigilance.

In the absence of salient information, an encounter with a conspecific creates uncertainty. Without prior knowledge that might indicate affiliative or antagonistic interaction, the organism is required to anticipate multiple potential outcomes^35^. Such unpredictability demands a specialized system for the rapid detection of sensory information, to ensure appropriate behavioural and physiological responses. The hypothalamic system is central to this process, integrating sensory inputs and regulating the neuroendocrine axis to coordinate adaptive responses to environmental challenges^15,16,36,37^. Within this framework, the rapid activation of the CRH^PVN^ neurons upon exposure to a social stimulus reveals a mechanism for immediate social threat evaluation. This functionality aligns with the broader role of hypothalamic circuits in predicting and responding to urgent physiological and environmental demands^13,38^. Such a predictive mechanism in the hypothalamus enables proactive regulation of energy and hydration. It therefore provides an adaptive advantage by enabling individuals to anticipate potential challenges before physiological imbalance or harm occurs^39–41^. This anticipatory control minimizes reaction time and ensures preparedness under uncertainty, a principle consistent with evolutionary models of risk management in which the cost of false alarms is far outweighed by the cost of failing to respond to genuine threats^42^. Extending this framework to the social context, our work highlights CRH^PVN^ neurons as a crucial hub for employing anticipatory mechanisms to facilitate the rapid evaluation of social threat. Given that CRH^PVN^ neurons regulate the HPA axis, this anticipatory signal likely primes physiological readiness^31^.

The behavioural significance of this rapid hypothalamic signal is evident in the structure of social investigation. Investigative behaviours, especially anogenital sniffing, are central to how rodents evaluate threat and the emotional state of conspecifics^14,29,30,43^. Our optogenetic inhibition experiments reveal that suppressing CRH^PVN^ activity during intruder exposure selectively diminished anogenital investigation toward threatening intruders, underscoring the necessity of this neuronal signal for targeted social assessment.

Moreover, with repeated exposures, we observed behavioural habituation without a corresponding reduction in CRH^PVN^ activity, a persistent neuronal vigilance consistent with the principle of asymmetric risk management and the maintenance of precautionary internal states ^42,44^. This dissociation suggests that while overt social behaviours adapt through experience, hypothalamic circuits maintain an internal representation of uncertainty, preserving readiness for unexpected aggression. Notably, novelty itself is an inherent component of both potential and realized threats, raising the question of whether CRH^PVN^ neurons are responding to novelty rather than threat per se. Our repeated-exposure experiments demonstrate that CRH^PVN^ neurons do not merely track novelty but instead act as uncertainty predictors, maintaining activity when the social environment retains an element of unpredictability.

Mounting evidence suggests that CRH^PVN^ neurons also signal centrally, either projecting directly to nearby nuclei or to long-range targets, allowing them to function at timescales that dynamically adjust the stress-response as threats evolve^17,45,46^. Neural projections rapidly influence stress-responsive behaviours, while CRH release at the median eminence elevates HPA axis activity and circulating cortisol levels, thereby coordinating immediate behavioural responses with longer-lasting endocrine adaptations to stress^36,47^. Further studies to identify the brain regions to which CRH^PVN^ neurons project, and the mechanisms by which they regulate this behaviour, will be relevant for understanding the central role of CRH^PVN^ neurons in rapid threat detection.

In summary, our findings identify CRH^PVN^ neurons as a rapid sensory-detection system that transforms social cues into adaptive behavioural responses. These neurons extend the predictive, feed-forward logic established for the homeostatic regulation into the social domain, linking the detection of potential threats with the preparation of appropriate physiological states. The discovery of a hypothalamic system that sustains social vigilance even as overt behaviour habituates provides a new framework for understanding how internal state and social perception interact. Because dysregulated CRH signalling and maladaptive threat evaluation are hallmarks of stress-related disorders, understanding this mechanism could inform the development of targeted strategies to restore balanced threat sensitivity and flexible social behaviour.

## METHODS

### Animals

All animal protocols were approved by the University of Calgary Animal Care and Use Committee (AC21-0067). Mice that were the offspring of *Crh-IRES-Cre* (B6(Cg)-*Crh*^*tm1(cre)Zjh*^; stock no: 012704) crossed with *Ai148 (Ai148(TIT2L-GC6f-ICL-tTA2)-D*; stock number 030328, Jackson Laboratories) were utilized for all optogenetics, fibre photometry and behavioural tracking experiments. Mice were group-housed on a 12:12 h light/dark schedule (lights on at 07:00 hours) with *ad libitum* access to food and water. CD-1 mice (Charles River Laboratory), aged over 3 months, were used for behavioural testing with an aggressive intruder. Mice were 6–8 weeks old at the time of surgery and virus injection.

### Viruses

AAV carrying Arch3.0-mCherry (LotAAV1081, AAV2/9-EF1a-DIO-Arch3.0-mCherry; Canadian Neurophotonics Platform Viral Vector Core Facility (RRID: SCR_016477) or mCherry (LotAAV1088, AAV2/9-EF1a-DIO-mCherry; Canadian Neurophotonics Platform Viral Vector Core Facility (RRID: SCR_016477)) was used for optogenetic manipulations.

### Stereotaxic injection and optical fibre implantation

Mice were maintained under isoflurane anesthesia in the stereotaxic apparatus. Mono fibre optic cannula with 400 µm diameter (Doric Lenses, MFC_400/430/0.48_5.5mm_MF2.5_FLT) was implanted dorsal to the PVN for fibre photometry (AP, −0.7 mm; L, −0.2 mm from the dura; DV, −4.85 mm from the dura) and for optogenetics (for Arch3.0: AP: −0.7 mm; L: 0.0 mm from the dura; DV: −4.5 mm) following tracking with a custom-made needle (AP: −0.7 mm; L: −0.2 mm from the dura; DV: −4.2 mm) and bilateral injections (AP: −0.7 mm; L: 0.2 mm from the dura; DV: −4.5 mm) fixed to the skull with METABOND® and dental cement. Mice were given 2 weeks to recover prior to the start of the experiment. Experiments started after an additional two weeks of recovery and handling.

### Histology

To verify GCaMP expression and ferrule location, following the experiments, mice were euthanized using isoflurane overdose and a subset of mice were transcardially perfused with phosphate-buffered saline (PBS), followed by 4% paraformaldehyde (PFA) in phosphate buffer (PB, 4 °C). Brains were placed in PFA 24 h followed by 20% sucrose PB. 50 µM coronal brain sections were obtained via cryostat. The other subset of mice was cut into 250 μm sections in ACSF, followed by fixation in 4% PFA, 1 M PBS, and 30% sucrose for 10 minutes each. Slide-mounted and coverslipped sections were imaged using a confocal microscope (Leica SP8). Images were prepared using ImageJ.

### Fibre photometry recording

Fibre photometry was used to record calcium transients from CRH neurons in the PVN of freely moving mice. After the recovery period, animals were handled for 5 min a day for 3 days and then habituated to the optic fibre in their home cage (5 min a day) for 3 days. We recorded 20 min of CRH^PVN^ neuron activity in the homecage immediately before and for 5 min after each test to improve bleaching correction. Doric fibre photometry system consisting of two excitation LEDs (465 nm and 405 nm from Doric) controlled by an LED driver and console, running Doric Studio software (Doric Lenses). The LEDs were modulated at 208.616 Hz (465 nm) and 572.205 Hz (405 nm), and the resulting signal was demodulated using lock-in amplification. Both LEDs were connected to a Doric Mini Cube filter set (FMC6_IE(400-410)_E1(460-490)_F1(500-540)_S)_E2(555-570)_F2(580-680)_S and the excitation light was directed to the animal via a mono fibre optic patch cord (DORIC MFP_400/430/1100-0.48_2m_FC/MF2.5). The LED power was adjusted to 30 µW at the end of the patch cord. The resulting signal was detected with a photoreceiver (Newport model 2151).

### Fibre photometry data analysis

Fluorescent signal data were acquired at a sampling rate of about 100Hz (Doric system). The data were then exported to MATLAB (MathWorks) for offline analysis using custom-written scripts (https://github.com/leomol/stimbox). Briefly, raw fluorescence traces were resampled to 20 Hz to reduce data size while retaining calcium signal dynamics. A 0.5 Hz low-pass filter was then applied to suppress high-frequency noise. The 465 nm and 405 nm data were first fit individually with a second-order polynomial, and the resulting fits were subtracted to remove bleaching artifacts. Next, a least-squares linear fit was applied to the 405 nm to align it with the 465 nm channel, and then the change in fluorescence (Δ*F*) was calculated by subtracting the 405 nm Ca^2+^ independent baseline signal from the 465 nm Ca^2+^ dependent signal at each time point. The output of these transformations yielded a motion-corrected dataset of changes in GCaMP6f fluorescence. For analysis, z-score calculation was performed using the following equation: ΔF/F = (F-F_0_)/ σF, where F is the test signal, F_0_ is the median signal of the baseline, and σF is the median absolute deviation of the signal.

### Optogenetics

Following the recovery period, mice were handled for 5 min on 3 consecutive days. Then, they were habituated to the fibre-optic cable attached to the cannula (without light) for 3 additional days (15-30 min per mouse).

For experiments, the light source (for Arch3.0: 561 nm, LRS-0561-GFO-00100-5, Laserglow Technologies) was connected to the Doric filter cube (FMC6_IE(400-410)_E1(460-490)_F1(500-540)_S)_E2(555-570)_F2(580-680)_S and the laser was directed to the animal via a mono fibre optic patch cord (DORIC MFP_400/430/1100-0.48_2m_FC/MF2.5). For Arch3.0, a yellow light (15 mW laser intensity) was used continuously for 20 seconds as the intruder was introduced to the resident’s home cage.

### Behavioural recordings and analysis

Adult male and female mice were used as residents. Resident mice encountered either a familiar or a novel intruder at each session. A familiar intruder is a conspecific from the same cage, whereas an unfamiliar intruder is from a different cage and strain (e.g., CD1 intruder). The test was conducted in the resident’s home cage. For video acquisition, a top-down camera (20 fps, Doric Studio) recorded all sessions under ∼30–50 lux of ambient light and was synchronized to the fibre photometry recording. These videos were later analyzed using the Behavioural Observation Research Interactive Software (BORIS, version 9^19^), and the following behaviours were scored: anogenital sniffing (directing the snout toward the anogenital area of conspecific); head sniffing (directing the snout toward the head of conspecific); torso sniffing (directing the snout toward the torso of conspecific); tail sniffing (directing the snout toward the tail of conspecific). Behaviour during 3 min after the introduction of the intruder was quantified and plotted. Quantification and analysis of behavioural ethograms, sliding time windows, cumulative probability, and transition probability were carried out using custom-written Python scripts.

### Statistics

Where quantification was made, data are represented as mean±standard error of the mean (s.e.m.). Statistical analysis was performed using GraphPad Prism 10.0. For comparisons, either two-tailed t-tests, ordinary or repeated-measures one-way ANOVA were applied as appropriate. When comparisons involved more than two groups and/or repeated factors, a two-way ANOVA was conducted, followed by Tukey’s or Sidak’s post hoc multiple-comparisons test. A significance threshold of p < 0.05 was applied for all analyses.

## ACKNOWLEDGEMENTS

We thank Mrs. Cheryl Breiteneder, Mrs. Dinara Baimoukhametova, Ms. Alexis Passmore, and Mr. Rodney Barasi for expert technical support. We also thank Dr. Matthew Hill for providing us with access to materials. We are grateful for the support of the Cumming School of Medicine Optogenetics Core Facility and the Hotchkiss Brain Institute Advanced Microscopy Platform.

This work was supported by an operating grant to J.S.B. from the Canadian Institutes of Health Research and the Brain Canada Neurophotonics Platform. I.A. was supported by the Canadian Institutes of Health Research-REDI-Early career transition award.

## AUTHOR CONTRIBUTIONS

I.A. and M.P. designed and conducted experiments, analyzed data, prepared figures, and wrote the paper. T.S designed experiments and contributed to the paper, T.F. designed experiments, discussed results and contributed to the paper. J.S.B. discussed results, prepared the paper and supervised the project.

