## Supplemental Figures for "Hypothalamic CRH neurons gate rapid social appraisal of conspecifics"

**Supplementary Figures**

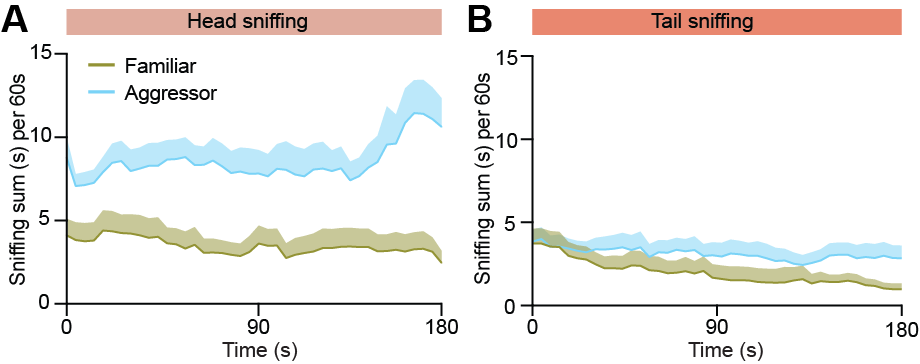

**FigureS1. Head and tail sniffing of a familiar versus aggressor intruder.**

**A.** Mean head sniffing time averaged in 60s bins with a 3s sliding window during the 3 min-long interaction (N = 12 for both groups – 2-way RM ANOVA Group X time: F (41,902) = 1.506, p = 0.022, shade represents s.e.m.). **B.** Mean tail sniffing time averaged in 60s bins with a 3s sliding window during the 3 min-long interaction (N = 12 for both groups – 2-way RM ANOVA Group X time: F (41,902) = 0.566, p = 0.987, shade represents s.e.m.).

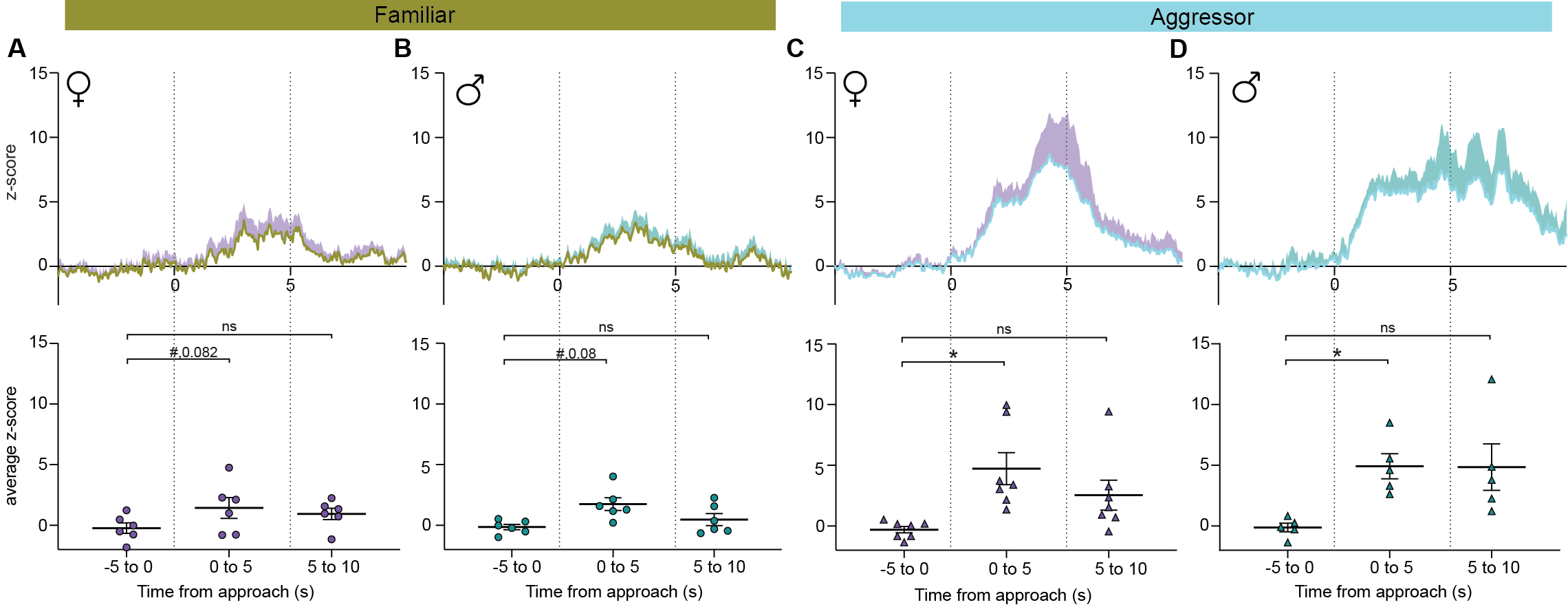

**Figure S2. Residents of both sex exhibit increased CRH^PVN^ activity towards an aggressor.**

**A.** Top: Group average trace of the z-scored calcium activity during social interaction of female residents towards familiar same-sex intruders. Bottom: change in z-score averages across the 5s time windows (w1,w2,w3), with 0s denoting approach corresponding to top panel (N=6 females, Repeated measures 1-way ANOVA, F (1.95,9.77) = 4.763, p = 0.036, Tukey’s multiple comparison test, w1 vs. w2 p =0.08, w1 vs w3 p = 0.16, w2 vs w3 p =0.65). **B.** Top: Group average trace of the z-scored calcium activity during social interaction of male residents towards familiar same-sex intruders. Bottom: change in z-score averages (N=5 females, Repeated measures 1-way ANOVA, F (1.93,9.65) = 4.88, p = 0.035, Tukey’s multiple comparison test, w1 vs. w2 p =0.08, w1 vs w3 p = 0.6, w2 vs w3 p =0.15). **C.** Top: Group average trace of the z-scored calcium activity during social interaction of female residents towards aggressor same-sex intruders. Bottom: change in z-score averages (N=7 females, Repeated measures 1-way ANOVA, F (1.7,10.21) = 11.09, p = 0.003, Tukey’s multiple comparison test, w1 vs. w2 p =0.014, w1 vs w3 p = 0.097, w2 vs w3 p =0.086). **D.** Top: Group average trace of the z-scored calcium activity during social interaction of male residents towards aggressor same-sex intruders. Bottom: change in z-score averages (N=5 females, Repeated measures 1-way ANOVA, F (1.44,5.77) = 7.98, p = 0.025, Tukey’s multiple comparison test, w1 vs. w2 p =0.027, w1 vs w3 p = 0.11, w2 vs w3 p =0.99).

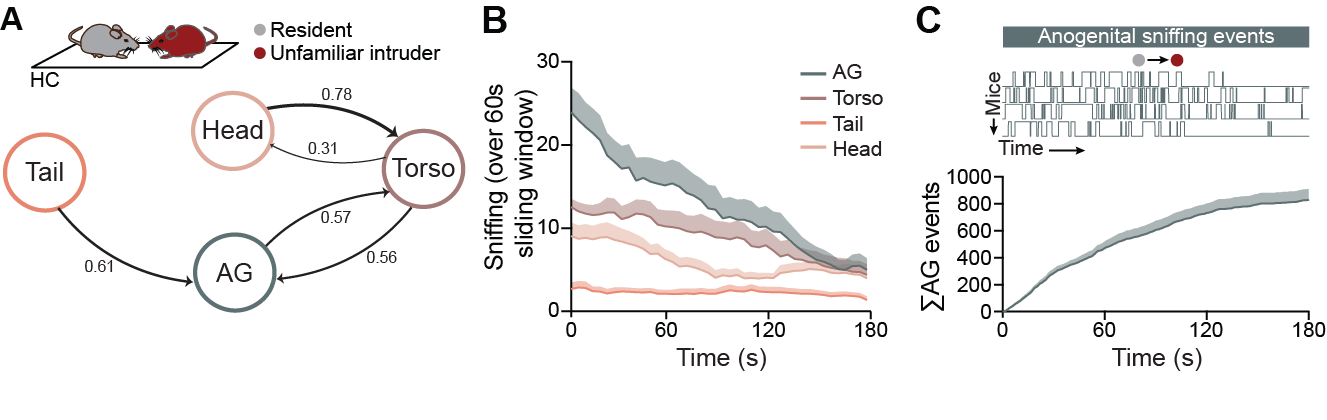

**Figure S3. Investigative behaviours of a resident towards an unfamiliar conspecific intruder.**

**A.** Transition probability plots representing switching of sniffing behaviours towards unfamiliar mice obtained from 12 mice across 3 min of interaction. **B.** Mean sniffing time averaged in 60s bins with a 3s sliding window, across different types of interactions (N=12). **C.** Example raster plots of anogenital sniffing events. **D.** Group average of cumulative duration for anogenital sniffing (N=12).

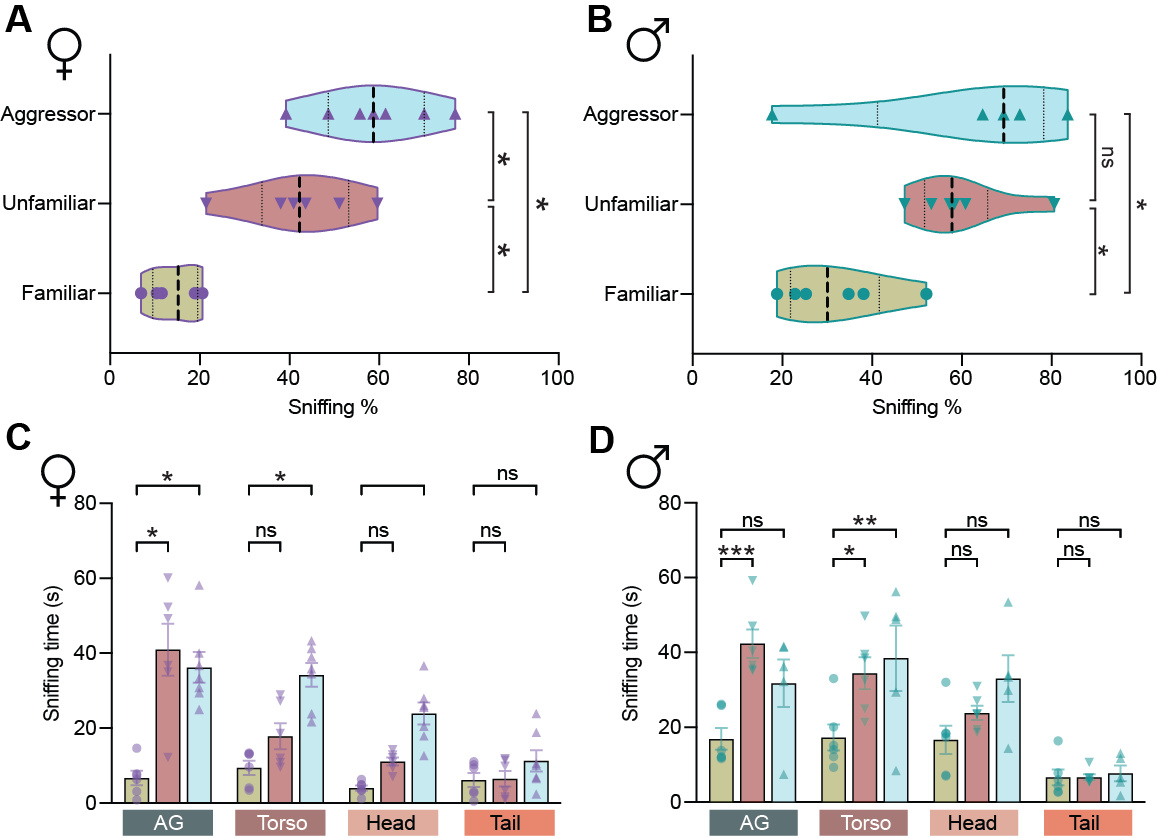

**Figure S4. Residents of both sex exhibit increased investigative behaviours towards threat-inducing intruders.**

**A.** Violin plots depicting the percentage of total sniffing by residents during 3-min-long social interactions of female residents towards the three types of same-sex intruders. (N = 6-7 per group, Ordinary 1-way ANOVA, F (2,16) = 18.6, p < 0.0001, Tukey’s multiple comparison test, Familiar vs. Aggressor p < 0.0001, Familiar vs. Unfamiliar p = 0.0013, and Unfamiliar vs. Aggressor p = 0.04). **B.** Violin plots depicting percentage of total sniffing by residents during 3-min long social interactions of male residents towards the three types of same-sex intruders. (N = 6-7 per group, Ordinary 1-way ANOVA, F (2,16) = 5.55, p < 0.016, Tukey’s multiple comparison test, Familiar vs. Aggressor p = 0.029, Familiar vs. Unfamiliar p = 0.033, and Unfamiliar vs. Aggressor p = 0.978). C.

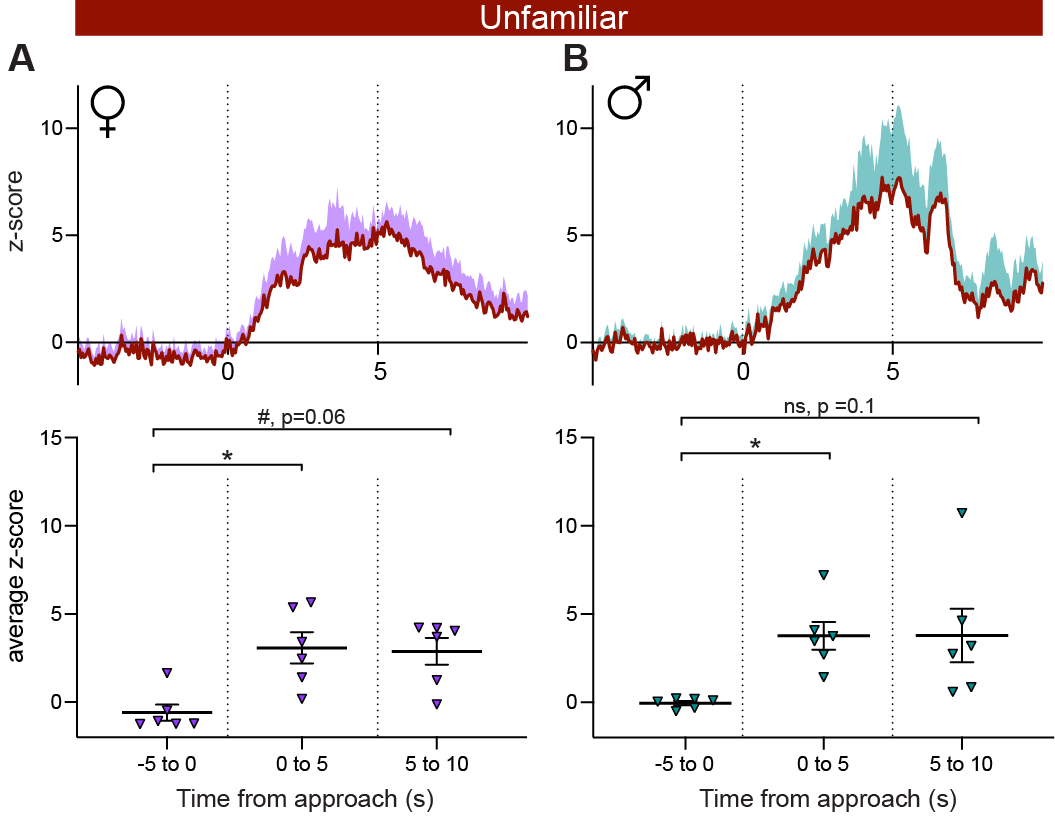

**Figure S5. Residents of both sex exhibit increased CRH^PVN^ activity towards an unfamiliar conspecific.**

**A.** Top: Group average trace of the z-scored calcium activity during social interaction of female residents towards unfamiliar same-sex intruders. Bottom: change in z-score averages across the 5s time windows (w1,w2,w3), with 0s denoting approach corresponding to top panel (N=6 females, Repeated measures 1-way ANOVA, F (1.5,7.8) = 10.22, p = 0.008, Tukey’s multiple comparison test, w1 vs. w2 p =0.011, w1 vs w3 p = 0.017, w2 vs w3 p =0.983). **B.** Top: Group average trace of the z-scored calcium activity during social interaction of male residents towards unfamiliar same-sex intruders. Bottom: change in z-score averages (N=5 females, Repeated measures 1-way ANOVA, F (1.05,5.29) = 8.25, p = 0.031, Tukey’s multiple comparison test, w1 vs. w2 p =0.011, w1 vs w3 p = 0.11, w2 vs w3 p =0.99).

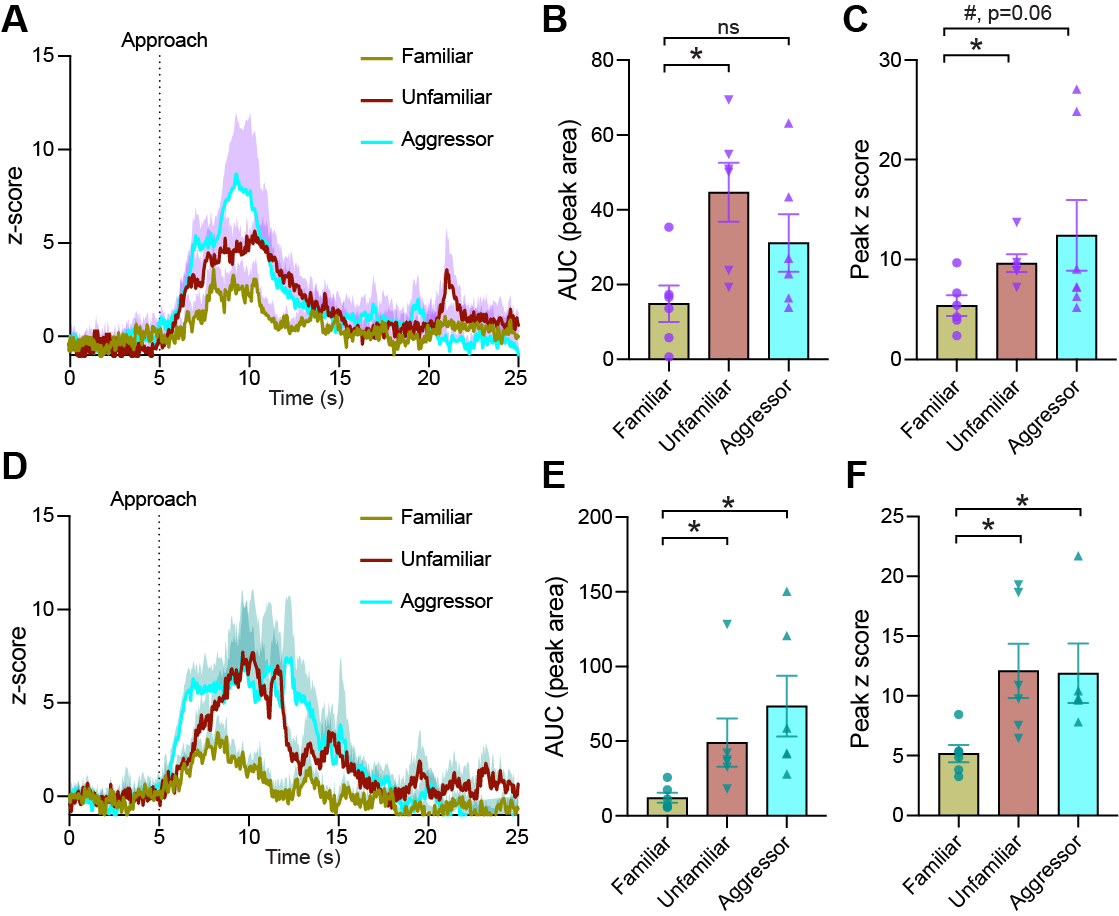

**Figure S6. CRH^PVN^ activity increases in both male and females during social appraisal of an intruder.**

**A.** Female averaged z-score of CRH^PVN^ calcium responses aligned at approach (shade represents s.e.m.). **B.** Individual peak area under the curve for the calcium activity for the first peak after approach in females (N = 6-7 per group, Ordinary 1-way ANOVA, F (2,15) = 4.61, p = 0.027). **C.** Individual peak z-scores of the first peak after approach in females (N = 6-7 per group, Ordinary 1-way ANOVA, F (2,33) = 6.08, p = 0.0056). **D.** Male averaged z-score of CRH^PVN^ calcium responses aligned at approach, (shade represents s.e.m.). **E.** Individual peak area under the curve for the calcium activity for the first peak after approach in males (N = 5-6 per group, Ordinary 1-way ANOVA, F (2,15) = 4.15, p = 0.036). **F.** Individual peak z-scores of the first peak after approach in males (N = 5-6 per group, Ordinary 1-way ANOVA, F (2,33) = 6.08, p = 0.035).

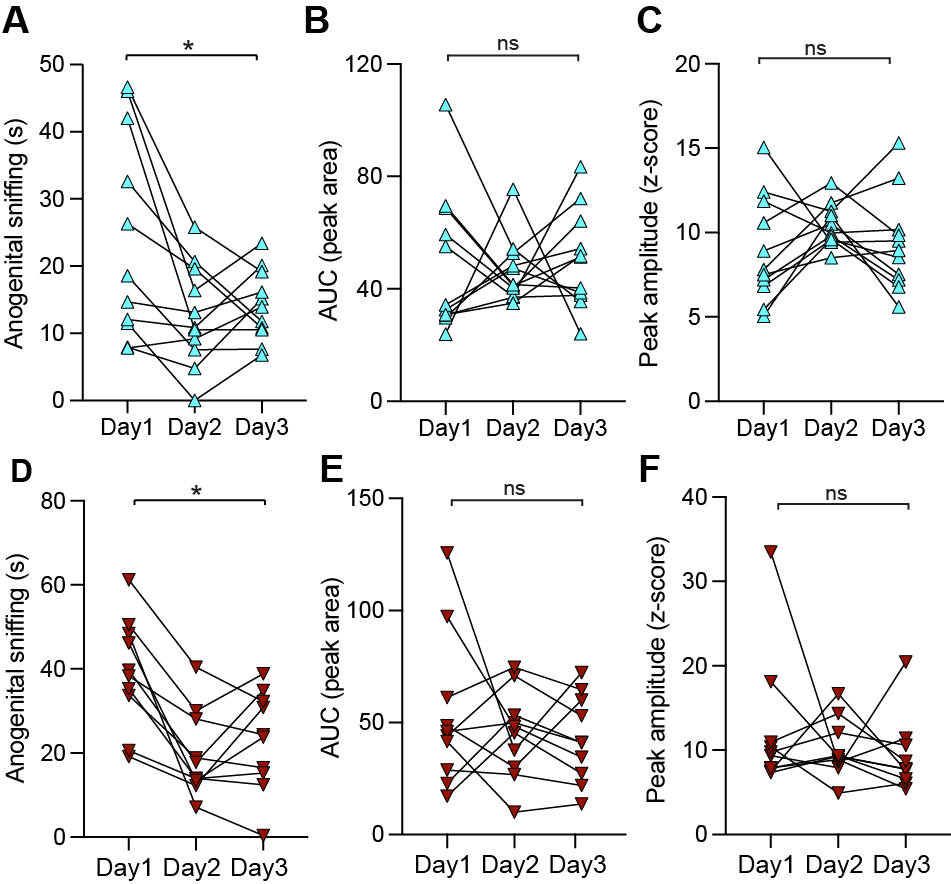

**Figure S7. Reduction in anogenital sniffing behaviour but not CRH^PVN^ activity upon repeated exposure of a threat-inducing intruder.**

**A.** Anogenital sniffing time of same aggressor intruder over 3 days (N=11, Repeated measures 1-way ANOVA, F (1.315,13.15) = 6.74, p = 0.016). **B.** AUC for CRH^PVN^ activity during approach to same aggressor intruder over 3 days (N=11, Repeated measures 1-way ANOVA, F (1.89,18.99) = 0.1, p = 0.88). **C.** Peak z-score for CRH^PVN^ activity after initiating approach to same aggressor intruder over 3 days (N=11, Repeated measures 1-way ANOVA, F (1.89,18.99) = 1.42, p = 0.26). **D.** Anogenital sniffing time of same unfamiliar intruder over 3 days (N=11, Repeated measures 1-way ANOVA, F (1.4,12.7) = 14.14, p = 0.0012). **E.** AUC for CRH^PVN^ activity during approach to same unfamiliar intruder over 3 days (N=11, Repeated measures 1-way ANOVA, F (1.29,11.63) = 0.68, p = 0.46). **F.** Peak z-score for CRH^PVN^ activity after initiating approach to same unfamiliar intruder over 3 days (N=11, Repeated measures 1-way ANOVA, F (1.59,14.36) = 0.75, p = 0.45).

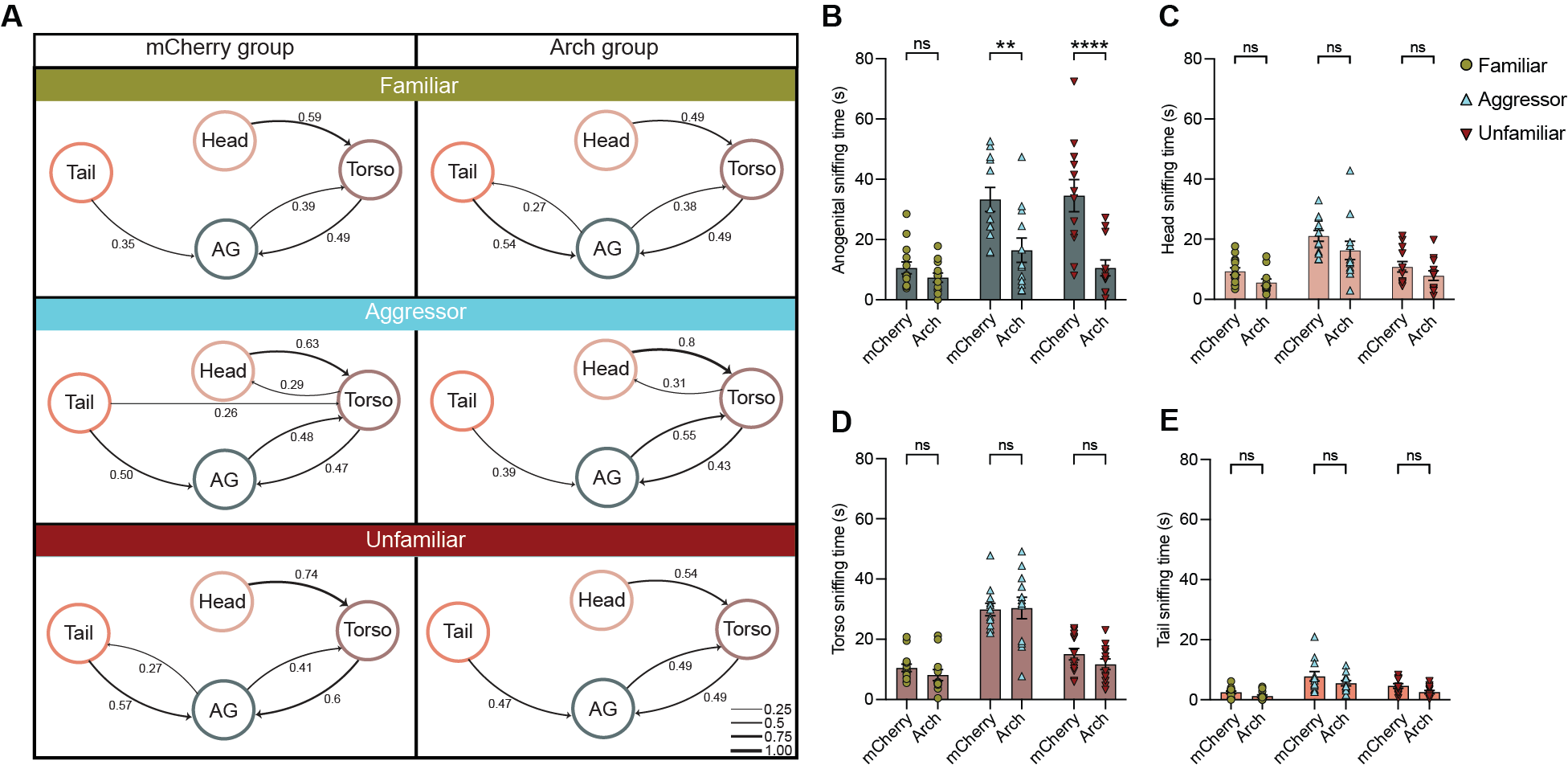

**Figure S8. Optogenetic inhibition of CRH^PVN^ activity during social appraisal results in targeted decrease in anogenital sniffing of an unfamiliar or aggressor intruder.**

**A.** Transition probability plots across three intruder types for optogenetic control and inhibited groups. **B.** Bar graph depicting mean ± s.e.m. of anogenital sniffing time for mcherry vs. arch residents towards all 3 types of intruders (N = 12-14 per group, Ordinary 2-Way ANOVA, F (2,68) = 4.77, p = 0.011, Sidak’s multiple comparison test, Familiar arch vs. Familiar mcherry p = 0.87, Aggressor arch vs. Aggressor mcherry p = 0.003, Unfamiliar arch vs. Unfamiliar mcherry p < 0.0001). **C.** Bar graph depicting mean ± s.e.m. of torso sniffing time for mcherry vs. arch residents towards all 3 types of intruders (N = 12-14 per group, Ordinary 2-Way ANOVA, F (2,68) = 0.42, p = 0.65, Sidak’s multiple comparison test, p > 0.5 for all comparison types on graph). **C.** Bar graph depicting mean ± s.e.m. of head sniffing time for mcherry vs. arch residents towards all 3 types of intruders (N = 12-14 per group, Ordinary 2-Way ANOVA, F (2,68) = 0.13, p = 0.87, Sidak’s multiple comparison test, p > 0.1 for all comparison types on graph). **D.** Bar graph depicting mean ± s.e.m. of tail sniffing time for mcherry vs. arch residents towards all 3 types of intruders (N = 12-14 per group, Ordinary 2-Way ANOVA, F (2,68) = 0.19, p = 0.81, Sidak’s multiple comparison test, p > 0.1 for all comparison types on graph).
