## Supplemental Tables for "Hypothalamic CRH neurons gate rapid social appraisal of conspecifics"

**Supplementary Table 1**

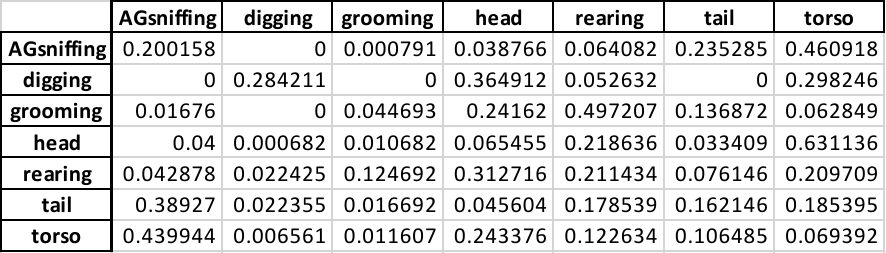

Table S1. Transition Probability values between behaviors for familiar group.

**Supplementary Table 2**

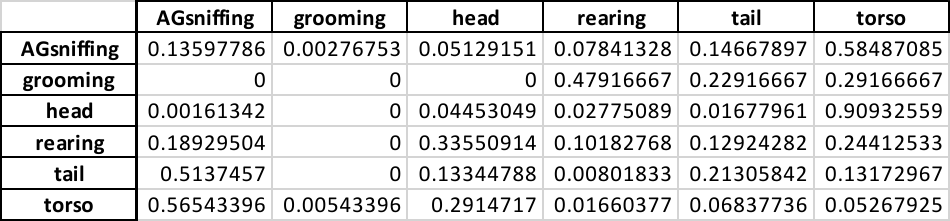

Table S2. Transition Probability values between behaviors for aggressor group.

**Supplementary Table 3**

|  | **AGsniffing** | **digging** | **grooming** | **head** | **rearing** | **tail** | **torso** |
| --- | --- | --- | --- | --- | --- | --- | --- |
| **AGsniffing** | 0.182704 | 0.000398 | 0.010007 | 0.01438 | 0.026508 | 0.195825 | 0.570179 |
| **digging** | 0 | 0.132653 | 0 | 0.214286 | 0.469388 | 0.183673 | 0 |
| **grooming** | 0 | 0 | 0.046083 | 0.105991 | 0.414747 | 0.036866 | 0.396313 |
| **head** | 0.083215 | 0.006697 | 0.005842 | 0.056569 | 0.053719 | 0.008264 | 0.785694 |
| **rearing** | 0.139956 | 0.016617 | 0.048744 | 0.302806 | 0.234121 | 0.090842 | 0.166913 |
| **tail** | 0.615957 | 0 | 0.018256 | 0.032454 | 0.145143 | 0.105477 | 0.082714 |
| **torso** | 0.563578 | 0 | 0.001981 | 0.314888 | 0.034569 | 0.049265 | 0.035719 |

Table S3. Transition Probability values between behaviors for unfamilar group.

**Supplementary Table 4**

| Type/Group | Familiar | Aggressor | Unfamiliar |
| --- | --- | --- | --- |
| AG | 28.2 | 31.9 | 45.9 |
| Torso | 31.9 | 33.4 | 28.8 |
| Tail | 15.2 | 9 | 6 |
| Head | 24.7 | 26.7 | 19.3 |

Table S4. Percentage occupancy of different types of sniffing in the pie chart in Figure 3D

**Supplementary Table 5**

| Type/Group | Familiar | | Aggressor | | Unfamiliar | |
| --- | --- | --- | --- | --- | --- | --- |
|  | mcherry | arch | mcherry | arch | mcherry | arch |
| AG | 32.2 | 32.8 | 36.3 | 23.9 | 53.1 | 32.3 |
| Torso | 31.9 | 36.5 | 32.1 | 44.3 | 23.1 | 35.8 |
| Tail | 7.5 | 5.4 | 8.5 | 8.1 | 7.2 | 7.7 |
| Head | 28.4 | 5.6 | 23.1 | 23.7 | 16.6 | 24.1 |

Table S5. Percentage occupancy of different types of sniffing in the pie chart in Figure 5D,I and N respectively.
